# Breeding behaviour and productivity of Black-necked crane (*Grus nigricolis*) in Ladakh, Indian Trans-Himalaya

**DOI:** 10.1101/809095

**Authors:** Pankaj Chandan, Tanveer Ahmed, Afifullah Khan

## Abstract

A long-term study was conducted to understand some aspects of breeding biology of Black-necked crane (*Grus nigricollis*) in Changthang, Ladakh. Data on aspects such as the breeding season, courtship, mating, egg laying and incubation period, nest site fidelity, egg morphometry, breeding productivity and recruitment rate were collected between 2003 and 2012. Black-necked crane started arriving from last week of March to first half of April and showed fidelity at ten nesting sites. Courtship and mating peaked early morning (0700 hours), around noon (1100 hours) and in late evening (0600 hours) while the nest building at evening (1600 hours). The egg laying period initiated in May and extended up to July. The average incubation period was 33.88±0.3 days. Hatching success, nest survival rate and fledgling increment rate of the bird were 73.3%, 0.55 ± 0.03 and 0.41 ± 0.02 respectively with an overall breeding productivity of 0.73. The present population of Black-necked crane in Changthang, Ladakh seems to stable with an average recruitment rate of 15.7±1.4.

## Introduction

The breeding of a species is largely influenced by the availability of food, water and temperature in boreal and temperate zones, while intensity of illumination and micro-habitat conditions apart from other demographic factors play an important role in tropics. The date at which each individual actually lays eggs is related to the food supply at the time of laying [1]. Laying cannot begin until food has become sufficiently abundant for the females needed for general body maintenance and for feeding their chicks. Most bird species breed around the time when food supplies are readily available and climatic conditions are favourable. In deserts, or in relatively well watered areas, the sexual cycles of vertebrates may be inhibited by drought and stimulated to sudden activity by rainfall [2].

For water birds like cranes, factors reported to influence nest success include, water depth at the nest [3-5], vegetation type around the nest [6-9], nest concealment [6], and land use practices [5]. A few studies have suggested influence of weather (e.g. annual precipitation) on the reproductive success [7]. The land use practices were reported to affect nest success [10] while predation is considered as one of the most important factors responsible for nest loss [11].

In Trans-Himalayan landscape, slowly melting glaciers and frozen wetlands mark the beginning of the biological activity for various water bird species including Black-necked cranes *Grus nigricolis*. However, available information on the breeding biology of black-necked crane in the Indian Changthang is scanty and largely based on convenience sampling, which is due to extreme and harsh climate of the area and limited funds and logistics. Except few long term studies [12-13], studies conducted so far on black-necked crane during the breeding season are mainly confined to explanatory notes on status at few sites and effects of dogs on the nesting birds [14-19]. Considering the paucity of scientific information on the breeding biology of black-necked crane, a long-term study to understand and collect information on aspects such as breeding season, courtship and display, nest site fidelity, egg laying, incubation and breeding productivity was conducted. There was also a need to ascertain recruitment rate of black-necked crane and factors influencing it, so that effective conservation and management strategy for long-term survival of the species can be suggested.

## Materials and Methods

### Study Area

Changthang Cold Desert Wildlife Sanctuary; henceforth Changthang (32°25’-34°35’ N & 77°30’-79°29’ E) declared in 1987 is located in the eastern part of Ladakh, Jammu and Kashmir state of India (Fig. 1). At an elevation between 3600 and 7000 m amsl., Changthang encompasses an area of about 22,588 km2. The Zanskar mountain Range forms its south-western border and the Karakoram Range the north-west. To the east, it is contiguous with Tibetan portion of Changthang, the westernmost extension of Tibetan plateau. The terrain is extremely rugged with a high proportion of open sandy plains, cliffs and exposed rocks. After Taklang La, the study area opens into an extensive plateau. The soil (pH ranges between 7 and 11), is sandy clay and to a lesser extent, loam or loamy clay [20-21], with poor organic matter and nitrogen content [22]. Extreme cold and arid conditions prevail in the area with monthly average maximum temperature during winters often remain less than 100 C while in summer it goes up as high as 270 C. The monthly average minimum temperature during winters fluctuates between sub-zero and −14 degree Celsius. Most of the precipitation is in the form of snowfall, and the average annual rainfall is about <100 mm.

**Fig 1.**
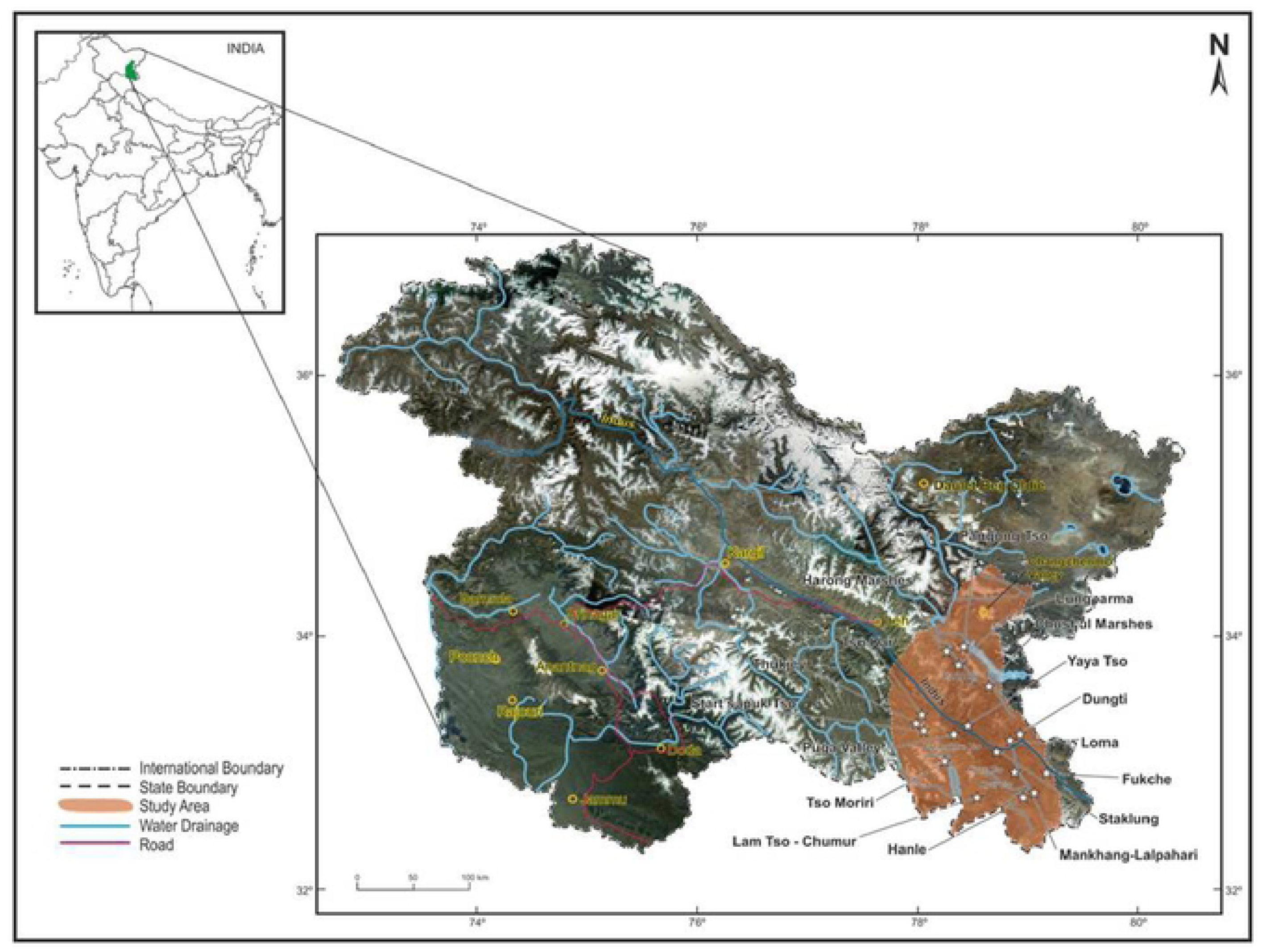
Location of Changthang cold desert Sanctuary in Ladakh, Jammu & Kashmir.

Changthang is virtually treeless, except for small isolated patches of Myricaria elegance and Yricaria germanica in addition to poplar (*Populus* spp.) and willow (*Salix* spp.) plantations at few places near human habitations. The major vegetation communities include *Caragana-Eurotia, Artemisia-Tanacetum, Stipa-Oxytropis-Alyssum* and *Carex melanantha-Leymus sccalinus*. At the higher altitudes (5000 m and above) sparse fell-field communities with moss or cushion-like growth forms, e.g. *Thylacospermum caespitosum, Arenaria bryophylla, Androsace saementosa* and lichens occupy the area. Changthang encompasses numerous brackish and freshwater wetlands; as small as one hectare to as large as 120 km^2^. Stream banks and marsh meadows around the wetlands exhibit characteristic sedge-dominated vegetation represented by species like *Carex, Kobresia, Scirpus, Triglochin, Pucciniella, Ranunculus* and *Polygonum* (Rawat & Adhikari, 2005). The shallow parts of the wetlands support a dense growth of aquatic plants such as *Hippuris vulgaris, Potamogeton pectinatus, Potamogeton perfoliatus, Zannichellia palustris*, and *Ranunculus natans*. These wetlands have been identified as the breeding grounds of several migratory and resident avian species including Black-necked Crane [18, 19, 23, 24].

### Methodology

To record breeding status of black-necked cranes’ population in Changthang, surveys were initiated in 2000, when two pairs of breeding cranes were seen, each at Tsomoriri and Tsokar basin. In subsequent years (2001 and 2002), extensive surveys were conducted covering large portion of Changthang plateau. All previously described sites were visited and occurrence of cranes was recorded at 21 different locations within Changthang. Once the distribution of cranes was understood, systematic surveys were initiated between 2003 and 2012 to collect information on various aspects of breeding biology.

Information on arrival and departure of Black-necked crane was collected from four different locations viz. Staklung, Chushul, Tsokar and Yaya Tso, since collection of data from all sites was not possible on weekly basis as the sites were widely distributed over an area of more than 20,000 Km^2^. These four sites, however, represent altitudinal and habitat variations between different breeding territories of Black-necked crane in Changthang. The initial and last sighting of Black-necked crane at any breeding ground was considered as arrival and departure dates respectively. Egg laying dates along with clutch size was also recorded each year. In addition to this information from locals and herders in the nearby villages around the nesting sites were also collected to ascertain the dates. Measurements of confirmed infertile eggs (34 eggs) were taken using a calliper.

Focal-animal sampling method (Altmann 1974) was used to determine the time budget of Black-necked crane during breeding season. Occurrence of breeding related behaviour such as courtship, mating and nesting were recorded at every five minute interval through a movable hide from morning (0600 hours) to late evening (1800 hours).

### Data Analysis

The egg volume was estimated using the following formula as described by (Hoyt 1979). Nesting success, hatching success, nestling survival rate, fledgling increment rate and overall breeding productivity were calculated to determine breeding success at various stages of breeding Black-necked cranes using the following formulae:

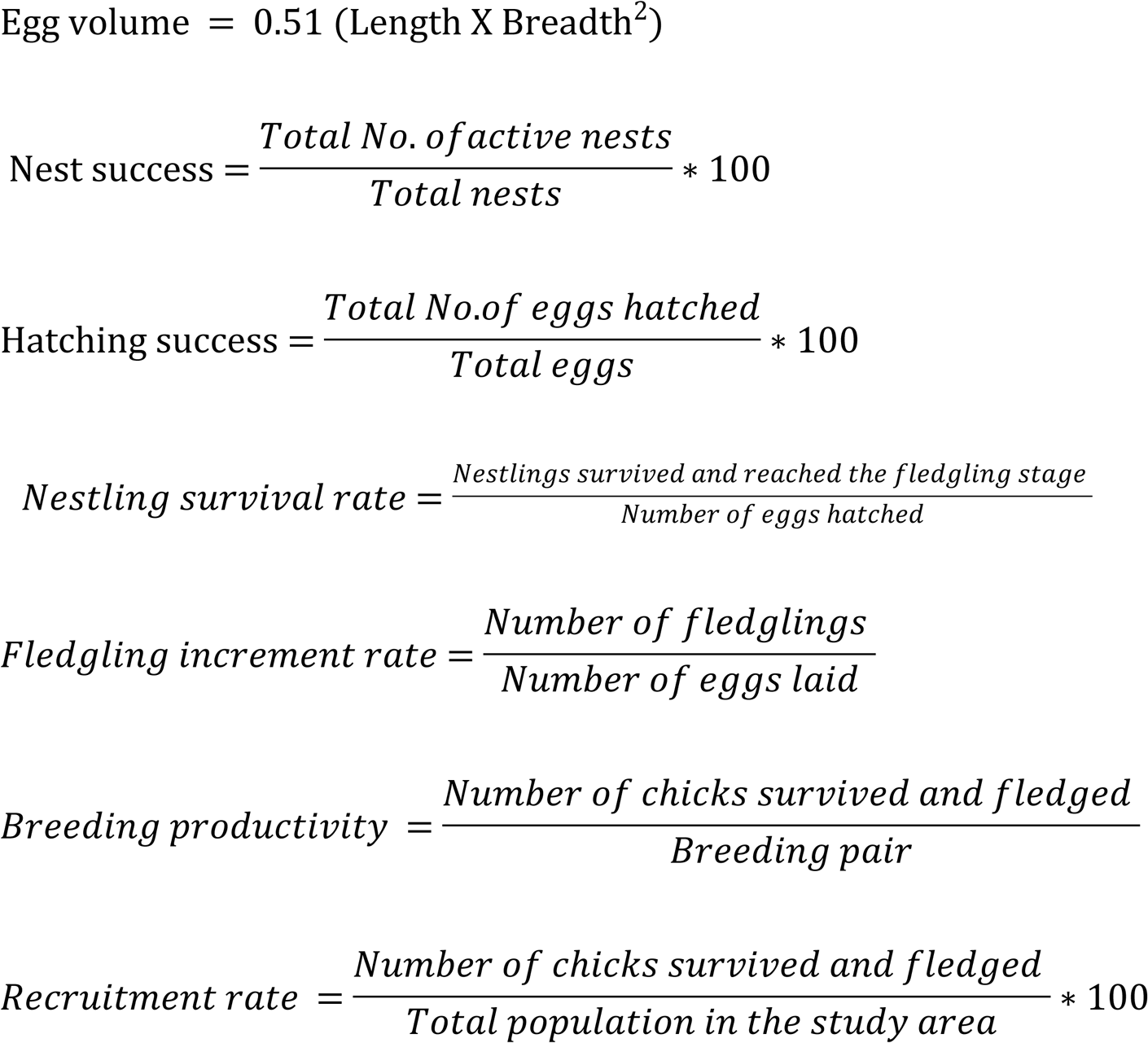

Data on courtship, mating and nest building were analysed by summarizing frequency of occurrences on hourly basis and percentages were depicted to understand the activity rhythm. Differences in mean incubation period across different years and sites were analyzed using Kruskal-Wallis One Way ANOVA. One sample t-test was used to check the differences in mean hatching success and nestling survival rates. All statistical analysis was conducted in SPSS 16.0.

## Results

### Breeding behaviour of Black-necked crane

The arrival period of Black-necked crane ranged from last week of March to first week of May and peaked during first half of April in Changthang, (Table S1). The courtship and mating activity started within two to three days of arrival of cranes in their respective breeding territories. Courtship and mating peaked early morning (0700 hours), around noon (1100 hours) and late evening (0600 hours) (Fig. 2). Nest building activity remained at its low in the morning hours and increased between 1000 and 1200 hours and peaked at evening (1600 hours) (Fig. 2). The species showed site fidelity at 10 nesting sites while at three sites, nesting occurred at different places. At the rest five sites, nesting did not occur regularly (Table S2).

**Figure 2:**
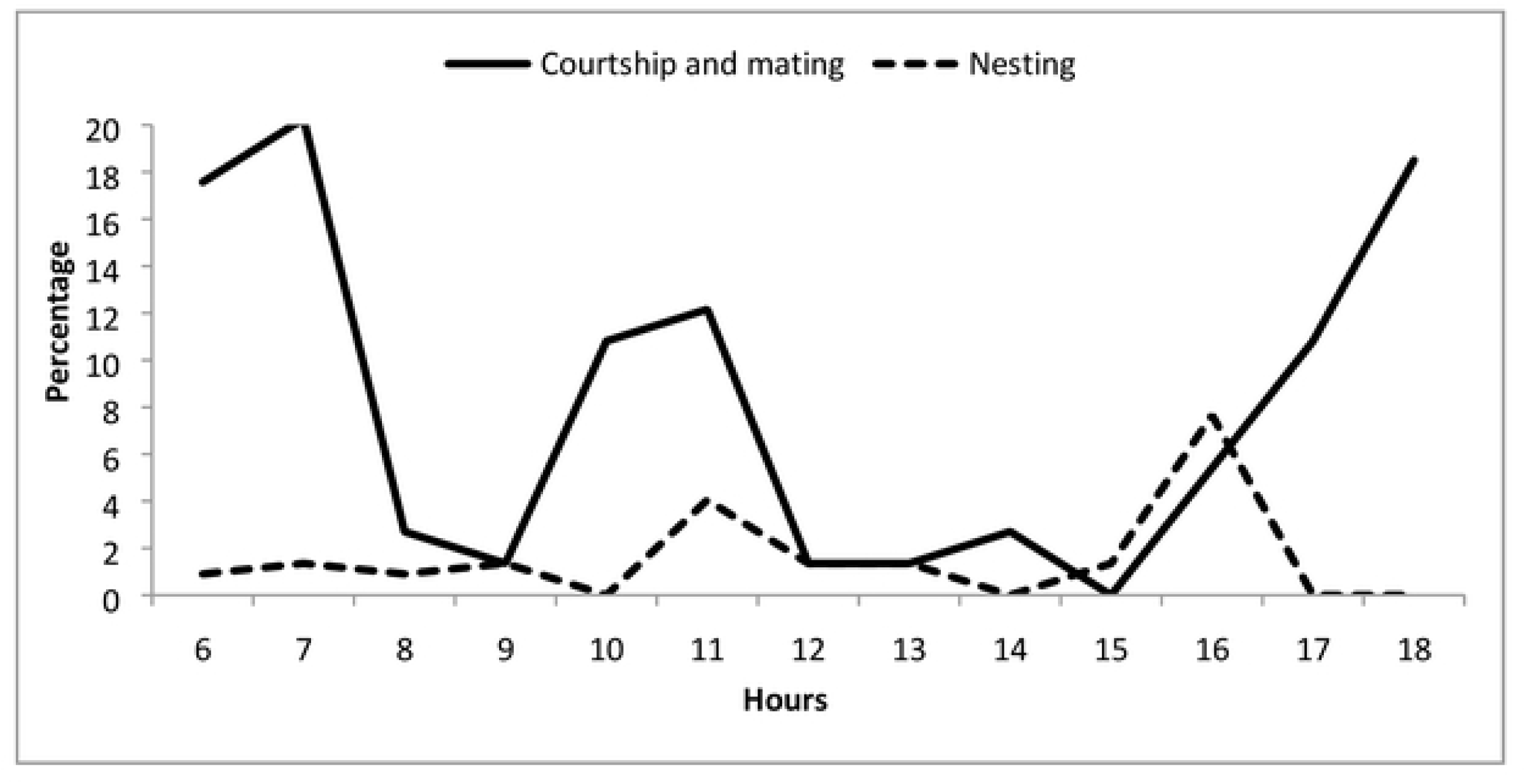
Diurnal rhythm of courtship-mating and nest building activity in Changthang, Ladakh.

A total of 146 nests of Black-necked crane were observed at 23 locations; 90.42% of nests were active while the rest (9.58%) nests were inactive. Of the total active nests, 76.03% were successful in raising at least one chick, while 23.97% of nests failed in raising any chick. The maximum nesting success (ca 86%) was recorded in 2005 and 2008 and a minimum (64.29%) was in 2012. The nesting success varied over different years of the study period (Fig. 3).

**Figure 3:**
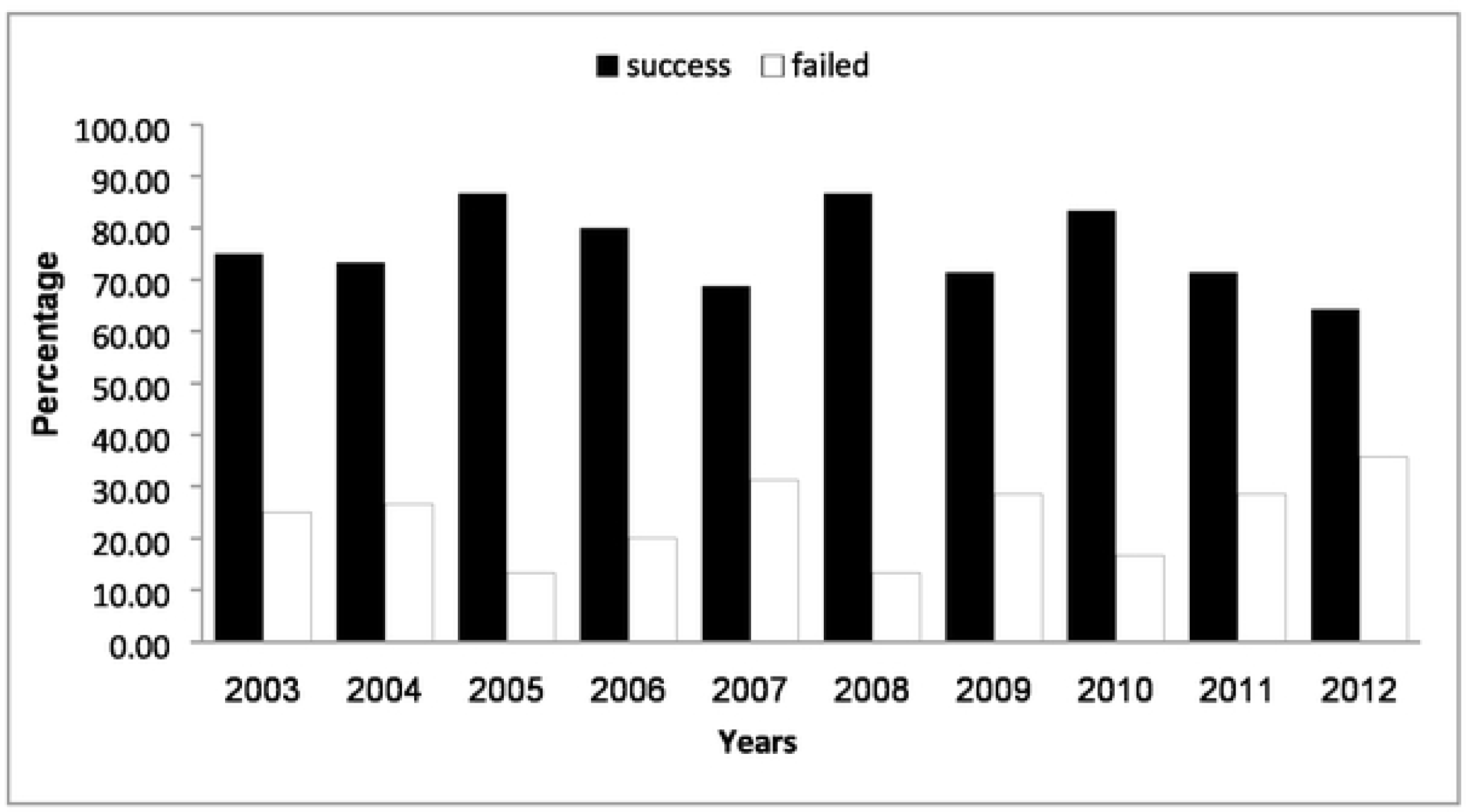
Percentage of successful and failed nesting during study period in Changthang, Ladakh.

**Figure 4:**
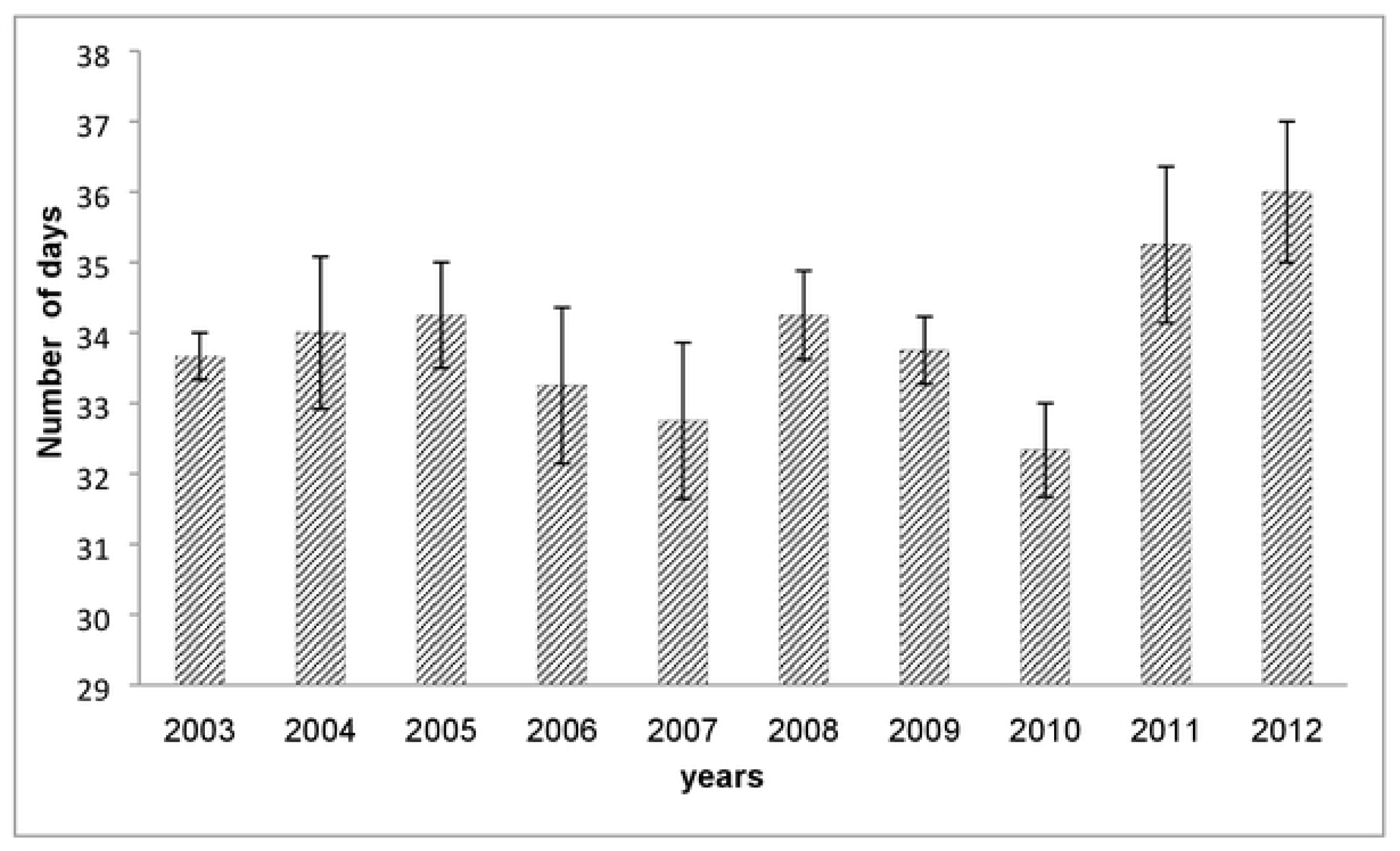
Average incubation period during different years in Changthang, Ladakh.

Most of the crane pairs laid their eggs during June (77.27%), nearly one fourth during May (19.09%) and few (3.64%) during July. The clutch size in most of the monitored nests (16 nesting sites) was two except one nesting site where single egg was recorded in 2005. The eggs of Black-necked crane had an average length breadth, weight and volume of 107.32 ± 1.02 mm, 63.17 ± 0.33 mm, 210.85 ± 3.17g and 218.95 ± 3.67 cm^3^ respectively (Table S3). The length, breadth, volume and weight of the eggs did not differ significantly over the years. Incubation started immediately after laying of the first egg with an average incubation period of 33.88 ± 0.3 days (range 30-38 days). The maximum (36.00 ± 1.0 days) incubation period was recorded during 2012 whereas minimum (32.33 ± 0.6 days) during 2010 (Fig. 3). The incubation period also did not differ significantly across years and various nesting sites.

### Breeding productivity of Black-necked crane

A total 266 eggs were laid during the study period, of which, 195 eggs (73.3%) were successfully hatched. The hatching success was maximum (92.9%) during 2008 and minimum (59.3%) during 2009 (Table 1). The hatching success of Black-necked crane differed significantly between different years of study (KW = 18.83, df = 9, p<0.01). Of the total eggs successfully hatched and produced nestlings, 108 chicks (56%) reached the fledgling stage. Nestling survival rate ranged between 0.44 and 0.68 with a mean value of 0.56 ± 0.03 (Table 1). The overall mean fledgling increment rate was 0.41 ± 0.03 (range 0.27-0.55). Overall breeding productivity of Black-necked crane was 0.74 while it was maximum during 2010 (0.92) and lowest (0.50) in 2011 (Table 1). The mean recruitment rate was 15.70±1.4% and it ranged between 7.91% and 22.03% during 2012 and 2006 respectively (Table 1).

**Table 1:** Breeding performance of Black-necked crane in Changthang, Ladakh

| Year | Total cranes | Breeding pairs | Hatching Success (%) | Nestling Success rate | Fledging Success rate | Breeding Productivity (%) | Recruitment rate (%) |
| --- | --- | --- | --- | --- | --- | --- | --- |
| 2003 | 60 | 16 | 73.3 | 0.45 | 0.33 | 0.63 | 16.67 |
| 2004 | 64 | 15 | 70.4 | 0.68 | 0.48 | 0.87 | 20.31 |
| 2005 | 58 | 15 | 76.7 | 0.43 | 0.33 | 0.67 | 17.24 |
| 2006 | 59 | 15 | 74.1 | 0.65 | 0.48 | 0.87 | 22.03 |
| 2007 | 58 | 16 | 66.7 | 0.61 | 0.41 | 0.69 | 18.97 |
| 2008 | 81 | 15 | 92.9 | 0.50 | 0.46 | 0.87 | 16.05 |
| 2009 | 65 | 15 | 59.3 | 0.56 | 0.33 | 0.60 | 13.85 |
| 2010 | 73 | 12 | 70.8 | 0.65 | 0.46 | 0.92 | 15.07 |
| 2011 | 79 | 14 | 61.5 | 0.44 | 0.27 | 0.50 | 8.86 |
| 2012 | 139 | 14 | 90.0 | 0.61 | 0.55 | 0.79 | 7.91 |
| Overall |  |  | **73.6±3.4** | **0.56±0.03** | **0.41±0.03** | **0.74** | **15.70±1.4** |

## Discussion

For many species of cranes, one of the most crucial unresolved questions related to breeding biology is one that concerns with breeding productivity over a longer period of time. An often asked research question while studying the breeding ecology of a species is to know the reproductive success of a nesting bird. Many consider clutch size, nesting success, and nesting mortality as critical site specific information required for long term conservation of the species. To bridge this information gap the present study has focused on some of these issues.

The Black-necked cranes start arriving in Ladakh during last week of March to the first week of May with majority arriving in the first half of April. Their departure starts during last week of October and continues until first week of November. However, Pfister [17] reported the arrival of Black-necked cranes on their breeding ground from late April to early May and departure from mid-October to November. The differences in the arrival and departure may be attributed to short term observations of Pfister [17], restricted only to one breeding season. Consistent with the present study, Dehao [25], Dwyer *et al*. [26] and Bishop [27] reported the arrival of Black-necked cranes on their breeding grounds in China from late March to mid-May and departure from mid-October to November. This similarity in the arrival and departure of cranes in India and China is due to the fact that Ladakh is part of the same eco-region. It is pertinent to note the late arrival of Black-necked cranes at Yaya Tso. Since Yaya Tso is the highest breeding place within Changthang, where the wetland remained frozen till about late April and hence crane arrive during last week of April and first week of May when melting of snow starts and weather is not so harsh.

The arrival was followed by courtship and mating which was frequently performed during the month of April and May. In most of the cases the courtship and mating was recorded during early morning, at about mid-day and in the evening. This is similar to the observation made during various other studies conducted on the breeding behaviour of Black-necked crane in China [26, 28-29].

The clutch size of most crane species is consistently two except grey-crowned and black-crowned cranes [30]. These two species of crane usually lay more than two eggs. The clutch of grey-crowned crane may even sometime consist of five eggs. However, such a variation has neither been observed not been reported from other breeding ranges of Black-necked crane. Larger clutch size of grey-crowned and black-crowned canes may be due to environmental conditions. Since these species inhabit tropical areas therefore they require less energy in maintaining body temperature as compared to Black-necked crane inhabiting alpine area. Furthermore, the former are resident while the letter is migratory. Most migratory species have small clutch size as they have to breed and rear chicks in relatively shorter span of time, before migration to their wintering areas.

Mean egg weight of Black-necked crane (210 g) observed during the present study was comparable as reported by Dehao *et al*. [25]. However, the mean egg weight during present study was of infertile eggs. We did not collect data from the active nests to avoid disturbance to the breeding pair and possible harm, if any, to the eggs. The incubation period of 30-38 days during present study was little more as reported by Dehao *et al*. [25] and Pfister [17]. The incubation period reported by the former was based on observation of three nests while the latter observed only one nest. During present study the sample size was large and therefore the variation was also large.

Hatching success of 27 avian species in the subarctic region ranged between 28 and 100 % and the major causes of hatching failure were attributed to weather conditions, predation and poaching [31]. The avian species with small clutch size had higher percentage of hatching success as compared to those had large clutch size [32]. Since most species of cranes have small clutch size it is therefore expected to have higher hatching success as compared to those avian species having large clutch size. The assertion seems to be true in case of crane species. The hatching success of at least 10 crane species other than Black-necked crane ranged between 56 and 93%; the lowest was reported for Grey crowned crane *Balearica regulorum* [33] and the highest was in Red-crowned crane *Grus japonensis* [34]. The hatching success observed during the current study is in conformity with that of the hatching species observed in other crane species. Predation, flooding, abandonment and egg infertility have been reported as major causes of hatching failure [26]. During present study average hatching success was 73.6%. The reasons of failure include depredation that accounted only about six per cent while two nests containing four eggs (about 1.3%) were washed away due to flood. Rest 20 % losses were for unknown reasons as it was not practically possible to continuously monitor all the nests, spread over such a large area as Changthang. Low hatching success in certain years could be attributed to weather condition.

High mortality of nestlings is probably a characteristic of majority of avian species and Black-necked crane is not an exception. Other species of cranes also show similar pattern of nestling survival. For instance, the fledgling percentage of conspecific Grey-crowned crane was 60 [33], Red-crowned crane 56 [35], Demoiselle crane *Grus virgo*, 63 [36] and it was 65 % for Florida sandhill crane *Antigone canadensis pratensis* [37]. The main cause of fledgling mortality in Changthang was depredation as nearly 25% fledglings were depredated by land and avian predators. The cause of mortality of rest 20% remained unknown.

A perusal of data on breeding productivity among different species of cranes revealed that it ranged between 0.13 and 1.05. The productivity of wattled crane *Bugeranus carunculatus* in Zambia was reported 0.13 [38]. It was 0.87 for common crane *Grus grus* [39], and 0.70 for sandhill crane *Antigone canadensis* [37]. The average breeding productivity (0.73) of Black-necked crane is more or less similar to other crane species. Of importance is the recruitment rate of a species in order to understand the status of population. During the present study, lowest recruitment rate observed during 2012 was due to sudden increase in wintering population of Black-necked crane in Changthang. The reasons of sudden surge in crane numbers were not known. Similar pattern was also observed in 2008 when the total population of wintering crane had increased from previous year’s 58 to 81. During 2011, the low recruitment is attributed to the egg predation and chick mortality. However, in rest of the years of study (excluding 2008 and 2012) the recruitment rate remained above 13%. Lovvorn and Kirkpatrick [40] opined that a stable breeding population of Sandhill cranes requires a recruitment rate of 10 to 12%. If the same criterion is applied, the present population of Black-necked crane in Changthang can be considered as stable.

## Acknowledgements

We are grateful to WWF Netherlands for long term funding support for this project. Thanks are due to Mr. Jigmet Takpa, CCF & Regional Wildlife Warden, Ladakh for granting permission to work in the area. A very special thanks to Mr. Ravi Singh, George Archibald, Ms. Esther Blom, Dr. Taej Mundukur, Mr. A. K. Singh, Dr. Sejal Worah, Dr. V. B. Mathur, Dr. Asad R. Rahmani, Maj. Gen. G. D. Bakshi, Mr. A. K. Srivastava, Dr. Parikshit Gautam, Dr. C. M. Seth, Ms. Archana Chatterjee, Dr. Dipankar Ghose, Late Parkash Gole, Col. R. T. Chacko, Prof. Jamal A. Khan, Dr. Gopi Sundar, Mr. Otto Pfister, Mr. Kiran Rajashekriah, Mr. Saleem-ul-Haq, Mr. Tsering Angchuk and Mr. Intesar Sohail for their help. We are also thankful to Dr. Li Fengshan, Mr. Blaise Humbert-Droz, Mrs. Usha Ganguli-Lachungpa, Mr. Shakeel Ahmed, Kamal Mehdi, Pushpendra Singh Jamwal, Rohit Ratan, Nisha Khatoon, Tsewang Rigzin, Anupam Anand, Pijush Kr. Dutta, Partha S. Ghose, Priyadarshinee Shrestha, Lak Tsheden Theengh, Mr. Ram Saroop and Mr. Dawa Tsering for their help and assistance.

## Supporting Information

Table S1: Arrival and departure dates of Black-necked crane at four study sites during various years in Changthang, Ladakh

Table S2: Nest sites fidelity at different breeding sites at Changthang, Ladakh

Table S3: Eggs morphometry of Black-necked crane in Changthang, Ladakh

